# Astrogliosis mapping in individual brains using multidimensional MRI

**DOI:** 10.1101/2022.01.10.475717

**Authors:** Dan Benjamini, David S Priemer, Daniel P Perl, David L Brody, Peter J Basser

## Abstract

There are currently no noninvasive imaging methods available for astrogliosis mapping in the central nervous system despite its essential role in the response to injury, disease, and infection. We have developed a machine learning-based multidimensional MRI framework that provides a signature of astrogliosis, distinguishing it from normative brain at the individual level. We investigated *ex vivo* cortical tissue specimen derived from subjects who sustained blast induced injuries, which resulted in scar-border forming astrogliosis without being accompanied by other types of neuropathology. By performing a combined postmortem radiology and histopathology correlation study we found that astrogliosis induces microstructural changes that are robustly detected using our framework, resulting in MRI neuropathology maps that are significantly and strongly correlated with co-registered histological images of increased glial fibrillary a cidic protein deposition. The demonstrated high spatial sensitivity in detecting reactive astrocytes at the individual level has great potential to significantly impact neuroimaging studies in diseases, injury, repair, and aging.

## Introduction

Astrocytes are glial cells that are spread throughout the mammalian central nervous system (CNS) where they represent the most abundant cell population in the brain [1]. As part of their many functions in the healthy CNS, astrocytes respond to CNS damage and disease through a process called astrogliosis, which occur in multiple CNS disorders including traumatic brain injury (TBI) [2, 3], autoimmune disease [4], stroke [5], neoplasia [6], and neurodegenerative diseases [7], and which plays an essential role in regulating CNS inflammation. The phenotypic cellular changes in astrocytes that are associated with astrogliosis can range from mild, with variable degrees of hypertrophy of cell body and stem processes, to that seen in scar-border forming astrogliosis, where cell processes overlap and intertwine to form compact borders [8]. The degree of glial fibrillary acidic protein (GFAP) deposition in reactive astrocytes often parallels the severity of the neuropathology [9] and is therefore the most widely used marker of astrogliosis.

Although reactive astrocytes are integral and essential components of CNS innate immunity and have numerous beneficial functions [10, 11], they can also cause harmful effects [2, 8, 12] that are regarded as detrimental to clinical outcomes. Regardless of the role astrogliosis plays in different conditions, it is a dominant feature and common component of almost all CNS disorders. However, the successful development of noninvasive imaging techniques, primarily ones based upon magnetic resonance imaging (MRI), to make astrogliosis visible has been elusive, mainly because of the failure of conventional MRI methods to detect it, but also, and perhaps more importantly, due to the experimental difficulty of disentangling astrogliosis from the neurological condition(s) that caused it. The latter is especially true in MRI and diffusion tensor imaging (DTI) studies involving TBI animal models that result in axonal injury, demyelination, neurodegeneration, edema, or neuroinflammatory processes that are concurrent with astrogliosis [13–16]. Studying astrogliosis is particularly difficult because of the challenges of decoupling the response to cellular alterations it generates from the response to the other microstructural and chemical processes that take place due to co-morbidities [17, 18].

In addition to the experimental difficulty of isolating astrogliosis in brain tissue, a basic limitation of MRI is its low spatial resolution – on the order of 1 mm^3^ on clinical scanners. Although relaxation and diffusion contrast mechanisms carry information about components at the micron length scale, coarse, voxel-averaged images “flatten” any intra-voxel heterogeneity, leading to loss of sensitivity and specificity in detecting microstructural changes induced by astrogliosis. There has been a recent push within the neuroimaging community to maximize the amount of information in an image by using a combination of magnetic field profiles to probe relaxation and diffusion mechanisms simultaneously, i.e., multidimensional MRI [19, 20]. That, combined with theoretical [21–24] and technological innovations [25, 26], have allowed the acquisition of MR images with effectively sub-voxel resolution, resulting in the identification of multiple biological components within a given voxel [27–31].

In this study we developed a multidimensional MRI machine learning framework to map astrogliosis in individual brains by focusing on blast induced TBI, which is a unique type of injury. Contrary to blast exposure, sequela of impact head injury (i.e., impact TBI) are well described [32]. Blast TBI is prevalent in the military cohort [33], and our understanding of the neuropathology following blast exposure is still in its infancy, particularly concerning its chronic sequelae. However, studies suggest that blast TBI is characterized by scar-border forming astrogliosis at brain interfaces including the subpial glial plate, around penetrating cortical blood vessels, at grey-white matter junctions (as illustrated in Fig. 1), and structures lining the ventricles, and which has been coined Interface Astroglial Scarring (IAS) [34, 35]. Importantly, scar-border forming astrogliosis is generally not accompanied by substantial axonal damage or phosphorylated tau (pTau) pathology, making these blast TBI cases ideally suited for studying chronic reactive astrocytes.

**Fig. 1.**
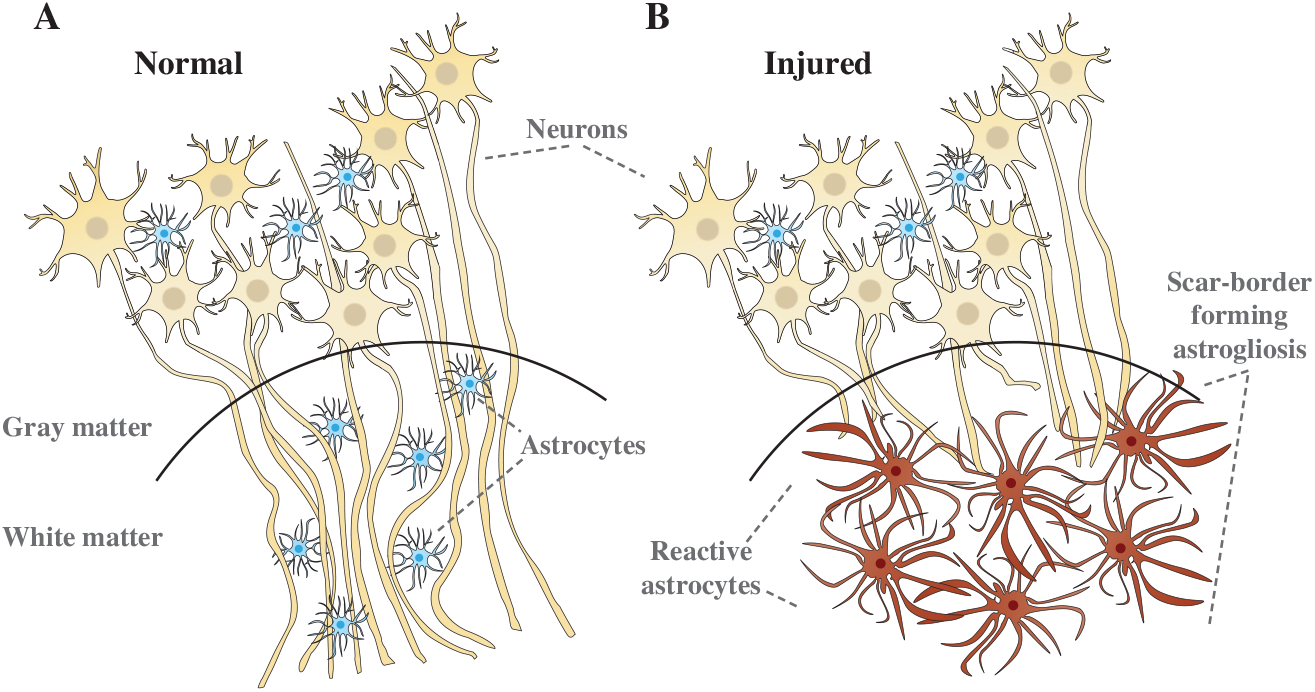
Illustration of microstructural changes occurring in the gray-white matter junction in (A) normal conditions, and (B) when scar-border forming astrogliosis is present. In (A) axons are tightly aligned forming a densely packed cellular environment. In (B) scar-border forming reactive astrocytes have overlapping processes that sequester damaged tissue and inflammation, while preventing injured axons from growing through the border. These changes are hypothesized to be reducing the overall cellular density in the white matter.

Here, we performed a combined postmortem multidimensional MRI and histopathology study to investigate the ways in which astrogliosis affects MRI relaxation and diffusion, and to establish whether multidimensional MRI can be used to map the presence of scar-border forming astrogliosis in brain tissue. We compared brain sections from blast-exposed military service members and from control individuals using robust quantitative radiological-pathological correlations, and developed a multidimensional MRI machine learning framework to map astrogliosis. We showed the spatial accuracy and sensitivity of the proposed framework, and its ability to produce results at the individual subject level. Our hope is that noninvasive mapping of astrogliosis would provide an important new tool for investigating and diagnosing a wide array of CNS disorders.

## Results

### Blast induced astrogliosis pathology

Table 1 and Supplementary Table 1 summarize the main demographic data and known medical history for each examined subject and histopathological findings observed in each studied tissue block. Of the total 14 cases evaluated, there were 7 cases with known IAS and 7 control cases negative for IAS.

**Table 1.** Main demographic and histopathological findings in patients with history of traumatic brain injury (TBI) and healthy controls

| Case<sup>†</sup> | Age | Manner of death | PMI (h) | Blast exposure | Impact TBI | Astrogliosis<sup>††</sup> |
| --- | --- | --- | --- | --- | --- | --- |
| 1 | 63 | Natural (cardiovascular) | 12 | None reported | None reported | None |
| 2 | 70 | Accident (MVA) | <12 | None reported | MVA | Mild |
| 3 | 60 | Accident (MVA) | N/A | None reported | MVA | Moderate |
| 4 | 48 | Natural (cardiovascular) | 22 | None reported | None reported | None |
| 5 | 52 | Suicide | N/A | None reported | Multiple concussions; MVA | None |
| 6 | 32 | Suicide | N/A | None reported | None reported | Mild |
| 7 | 59 | Suicide | 21 | None reported | Multiple MVAs | None |
| 8 | 38 | Suicide | N/A | IED exposure with LOC | As secondary injury | Scar-forming |
| 9 | 46 | Suicide | N/A | Multiple | Multiple concussions; MVA | Scar-forming |
| 10 | 29 | Suicide | <48 | Multiple | Fall with LOC | Scar-forming |
| 11 | 35 | Undetermined | <56 | Multiple | None reported | Scar-forming |
| 12 | 40 | Suicide | N/A | At least one exposure | Multiple concussions | Scar-forming |
| 13 | 64 | Natural (cardiovascular) | <48 | None reported | None reported | Scar-forming |
| 14 | 44 | Suicide | N/A | Multiple | Multiple MVAs | Scar-forming |

Postmortem interval (PMI); Traumatic brain injury (TBI); Motor vehicle accident (MVA); Improvised explosive device (IED); Loss of consciousness (LOC).
<sup>†</sup>All subjects in this study were males
<sup>††</sup>Microscopically confirmed GFAP-positive

Scar-border forming astrogliosis pathology is demonstrated in immunostained sections for GFAP from four representative cases in Fig. 2. Consistent with previous findings [34], blast-exposed subjects were characterized by dense astrogliosis at brain interfaces, including the grey-white matter junctions. The astrogliosis pathology in our cohort was notably present at the gray-white matter junction in white matter (WM), without associated accumulation of pTau in involved cortical regions, and otherwise not seen in the pattern of a known tauopathy (e.g., chronic traumatic encephalopathy, CTE). Astrogliosis did not coexist with axonal injury that would be indicated by amyloid precursor protein (APP) immunohistochemistry (Supplementary Figure 1). Additionally, hematoxylin and eosin (H&E)-based staining did not reveal evidence of ischemic-necrotic lesions, presence of vascular lesions, microhemorrhages, or tissue rarefaction.

**Fig. 2.**
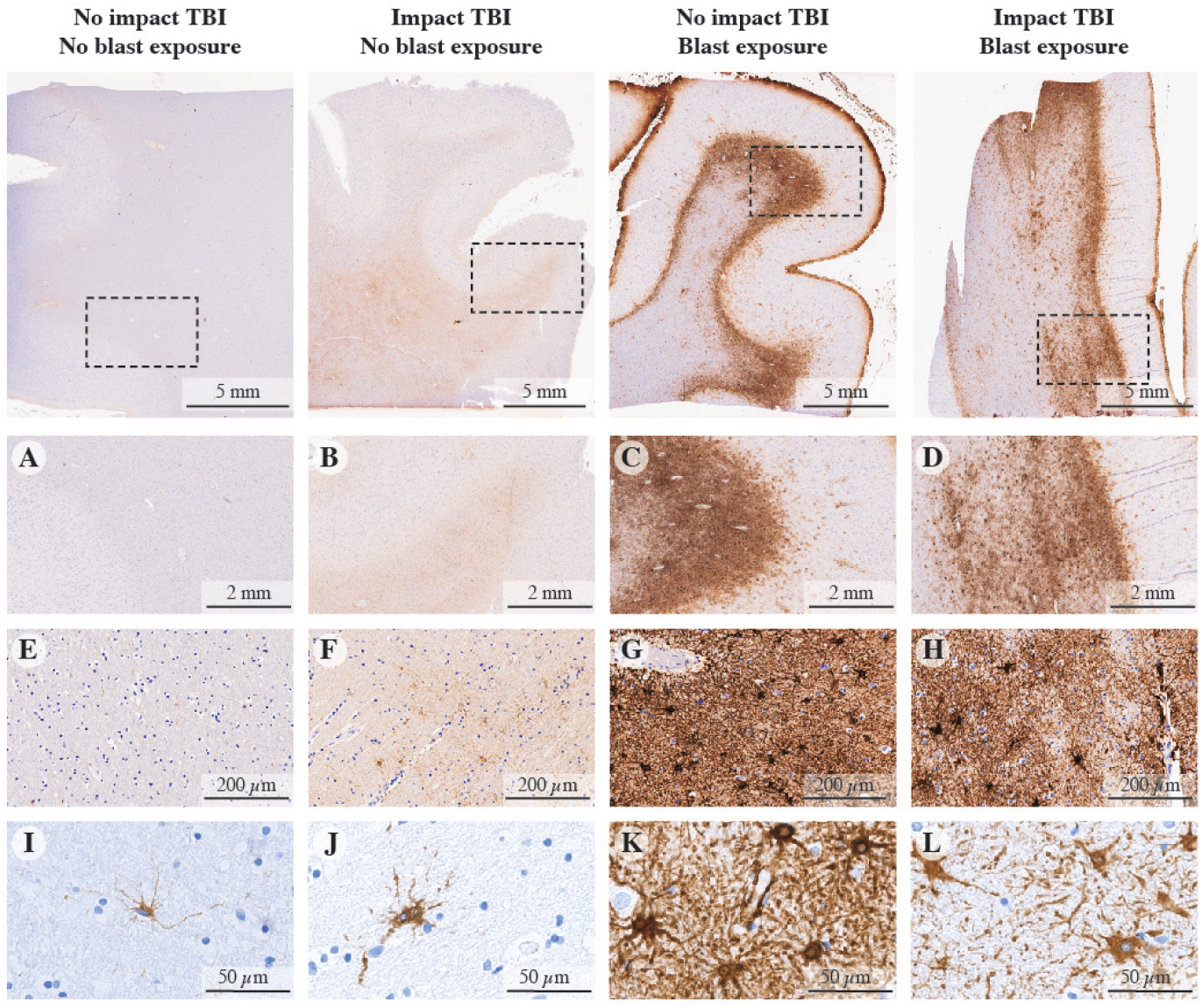
GFAP immunoreactivity in specimens without impact or blast exposure TBI, with impact TBI but without blast exposure, without impact TBI but with blast exposure, and with both impact and blast exposure TBI cases. (A), (E), and (I) show minimal GFAP immunoreactivity (Case 1). (B), (F), and (J) show limited GFAP immunoreactivity with mild reactive astrocytes (Case 3). (C), (G), and (K) show dense scar-border forming astrogliosis at the grey–white matter junction (Case 11). (D), (H), and (L) show a similar pattern of dense scar-border forming astrogliosis at the grey–white matter junction (Case 8).

There was a single IAS-positive case in which there is no reported history of blast or impact TBI exposure (Case 13); the cause of the pathology in this case is uncertain and may relate to under-reported TBI history.

### Astrogliosis has a multidimensional MRI signature

Our multidimensional MRI data reveal that a distinct signature exists for scar-border forming astrogliosis, which cannot be seen using one-dimensional MRI measurements. Investigation of the spatially-resolved subvoxel multidimensional spectral components illustrates these findings, and to this end, it is useful to summarize the rich dataset in a visually accessible manner. Each image slice of this data contains 4D information consisting of spatially-resolved spectra with 50*×*50 elements in each voxel. We can visualize these data as arrays of maps with varying subvoxel *T*_1_, *T*_2_, and mean diffusivity (MD) values. To make them more readable, the 50*×*50 spectra were sub-sampled on a 16×16 grid. Such summarized data of the *T*_2_-MD contrast from representative control (Case 7) and injured (Case 10) subjects are shown in Figs. 3 A and B, respectively. In addition, the marginal distributions (i.e., 1D spectra) of subvoxel MD values (top row) and subvoxel *T*_2_ values (right column) are shown to illustrate the information content of any 1D approach (yellow frames in Figs. 3 A and B). The GFAP histological image of each case is also shown on the upper-left corner of each panel, for reference.

**Fig. 3.**
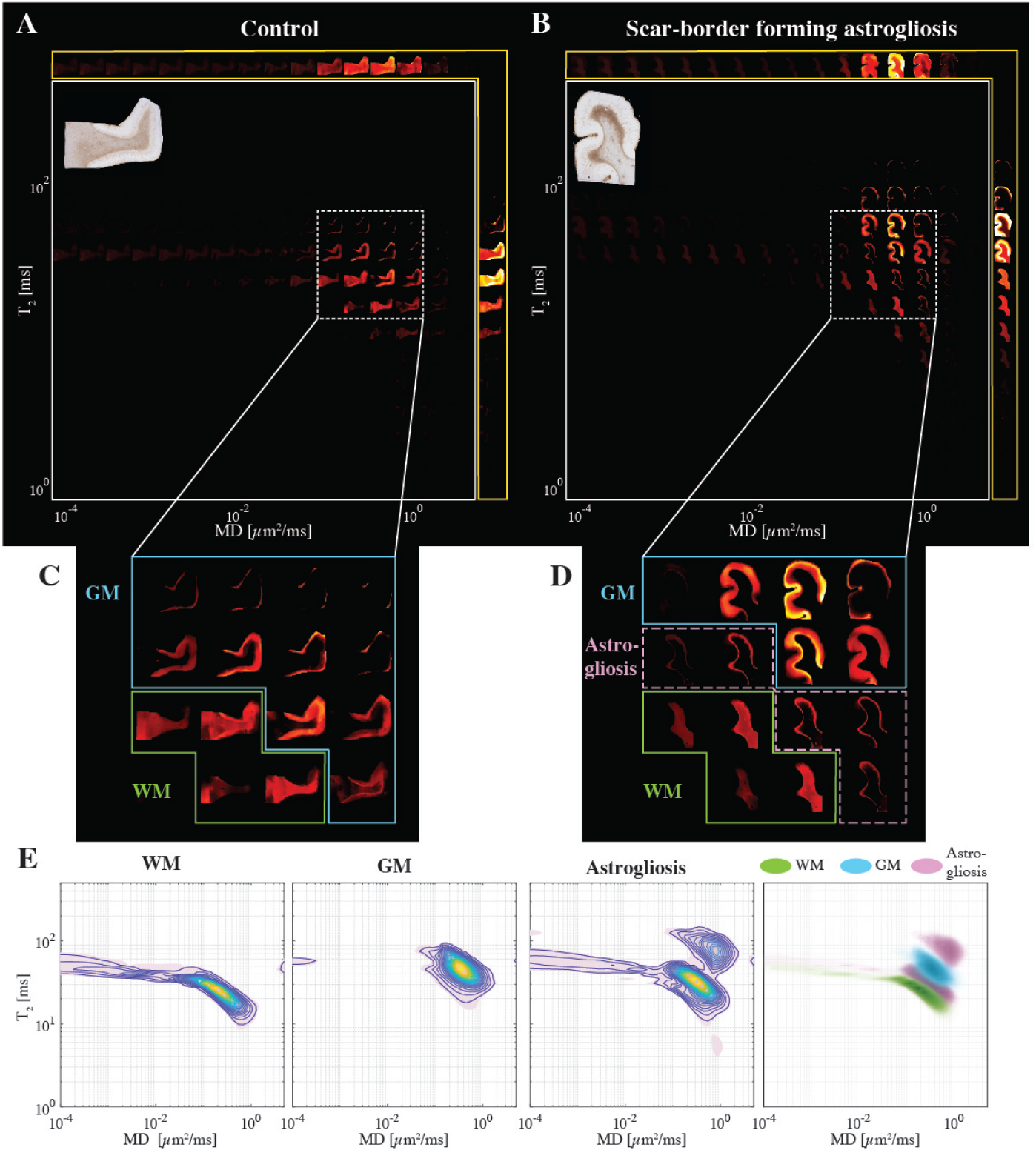
Changes in the *T*_2_-MD multidimensional MR signature induced by confirmed astrogliosis. Maps of 2D spectra of subvoxel *T*_2_-MD values reconstructed on a 16 16 grid of a representative (A) control (case 7) and (B) injured (case 10) subjects, along with their respective GFAP histological image (top left of each panel). (C) Magnified spectral region from the control case shows the clear separation of white (yellow frame) and gray (teal frame) matter according to their diffusion and *T*_2_ values. (D) The same magnified spectral region as in (C) from the injured case shows that while the WM and GM spectral information content is still clearly separable (yellow and teal frames, respectively), new WM-specific spectral components can be seen on the gray-white matter interface (purple frame), which is qualitatively similar to the GFAP staining pattern of the sample. (E) *T*_2_-MD spectra averaged across all subjects in WM, GM, and GFAP-positive spatial regions of interest (ROIs, left to right), and a superposition of the average spectra from the three ROIs.

The scar-border forming astrogliosis multidimensional MRI signature can be seen by examining the *T*_2_-MD range that contains most of the spectral information, highlighted as white rectangles. Magnifications of this range in the spectra are shown below in Figs. 3 C and D for the control and injured subjects, respectively. A clear separation of gray (blue frame) and white matter (green frame) can be seen in both control and injury states. However, we identified a distinct diagonal *T*_2_-MD spectral region (pink frame, Fig. 3D) in which intensities are concentrated at the gray-white matter junction on the WM side; these intensities follow closely the GFAP histological pattern (see inset image in Fig. 3B), while this newly found spectral information is absent in the control subject (Fig. 3C). Furthermore, the diagonal pattern in *T*_2_-MD points directly at a joint dependency with respect to *T*_2_ and MD, making it clear that this unique injury-related information cannot be seen by looking at *T*_2_ or MD separately (i.e., conventional 1D relaxation or diffusion MRI).

Generalizing these findings, we move from representative cases to averaged normal-appearing WM, gray matter (GM), and astrogliosis *T*_2_-MD spectra across the entire study (Fig. 3E left to right, respectively). A visualization of these three spectral clusters together is shown in the outermost right panel in Fig. 3E. The clear GM-WM separation can be easily seen, and in addition, the astrogliosis-related spectral information can be seen with most intensities lying in between the GM and WM peaks (gray-white matter junction), and some residual intensities towards higher values of *T*_2_ and MD (GM and meningeal border).

We also examined the *T*_1_-MD and *T*_1_-*T*_2_ datasets. We found that while they contain astrogliosis-related information, it is significantly reduced and harder to distinguish as compared with the *T*_2_-MD contrast, thus the latter provides the most potentially useful information. Nevertheless, summarized *T*_1_-MD and *T*_1_-*T*_2_ data are shown in Supplementary Figures 2 and 3, respectively.

### Anomaly detection in individuals

Although we demonstrated in the previous section that a multidimensional MRI signature associated with astrogliosis exists, being able to detect and refine it in an unsupervised and objective manner presents a significant challenge, especially when the information is hard to discern (e.g., *T*_1_-MD and *T*_1_-*T*_2_ in Supplementary Figures 2 and 3). In this work we build on previous frameworks [31, 36] and propose a within-subject anomaly detection procedure that results in MRI neuropathology biomarker maps. Conceptually, the principle is to first define what the ‘normal’ (i.e., uninjured in the desired context) multidimensional MRI signature is for a given individual, and then look for deviations from that ‘normal’. Here we implemented this framework by integrating co-registered histological images with multidimensional MRI data by using the GFAP density to define what is normative. Alternatively, the normative brain multidimensional signature could eventually be defined by collecting baseline multidimensional MRI data from healthy participants *in vivo*.

A schematic representation of the pipeline is shown in Fig. 4, where after the co-registration and deconvolution steps (Fig. 4 A-C), the GFAP density image is inverted to obtain a ‘normal’ mask in the image domain (Fig. 4 D). This mask is then applied on the multidimensional MRI dataset to isolate voxels outside of the injury regions. Once all normal-appearing voxels are identified within a subject, a Monte Carlo cross-validation procedure [37] is used to create multiple random splits (*N_cv_* = 1, 000 in this study) of 66% and 34% of the normal-appearing voxels into training and validation data, respectively. For each such split, the average multidimensional signature is computed using the training data (Fig. 4F), then thresholded to obtain a binary spectrum (i.e., spectral mask) of the normal-appearing tissue (Fig. 4G). In the next step, the inverse of this spectral mask is assumed to contain abnormal spectral information and is used, after binary dilation, voxelwise on the full multidimensional data to result in a map of abnormal signal components. This process is repeated *N_cv_* = 1, 000 times to allow the assessment of uncertainty and predictive accuracy, resulting in a set of neuropathology-related maps (Fig. 4H). The results are then averaged over the splits, yielding the final neuropathology MRI biomarker map (Fig. 4I).

**Fig. 4.**
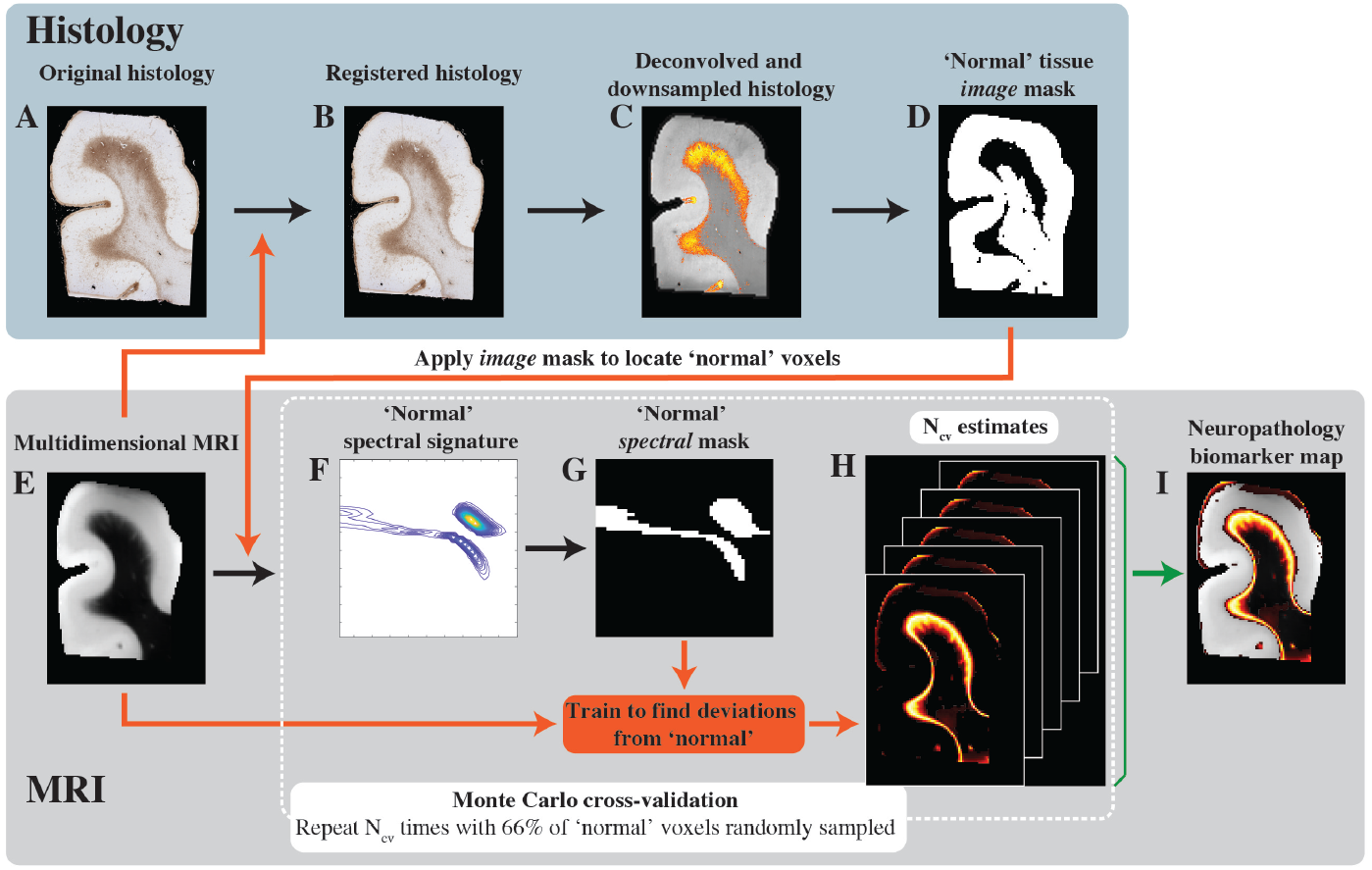
Schematic representation of the proposed anomaly detection framework. (A) The original GFAP histological image is processed in two steps: (B) co-registration to the MRI dataset, and (C) subsequent deconvolution and downsampling to match the MRI resolution. (D) This GFAP density image is then thresholded, inverted, and used as an image domain mask for normative brain voxels on the (E) multidimensional MRI data. A Monte Carlo cross-validation procedure is used to create *Ncv* = 1, 000 multiple random splits of 66% and 34% of the normal-appearing voxels into training and validation data, respectively, resulting in a 1,000 (F) normative spectral signatures, each of which is binarized to obtain spectral masks of normative brain. To detect anomalies, the normative spectral mask is inverted and is used on the full multidimensional data to directly obtain (H) *Ncv* = 1, 000 versions of abnormal signal components maps, which are then averaged to yield the final (I) neuropathology MRI biomarker map.

This strategy to detect anomaly in individuals was used separately on each subject and with each of the *T*_2_-MD, *T*_1_-MD, and *T*_1_-*T*_2_ datasets. It should be noted that it can be applied to any multidimensional data, with any number of dimensions.

### Multidimensional MRI maps of astrogliosis

First, we examined the performance of our machine learning framework in visualizing astrogliosis by assessing their spatial sensitivity and specificity. Figure 5 shows conventional MRI and DTI maps, multidimensional MRI maps, and histological GFAP density images of six representative control and injured cases (Cases 3, 4, 7, 10, 11, 12 shown in Figs. 5 A-F, respectively).

**Fig. 5.**
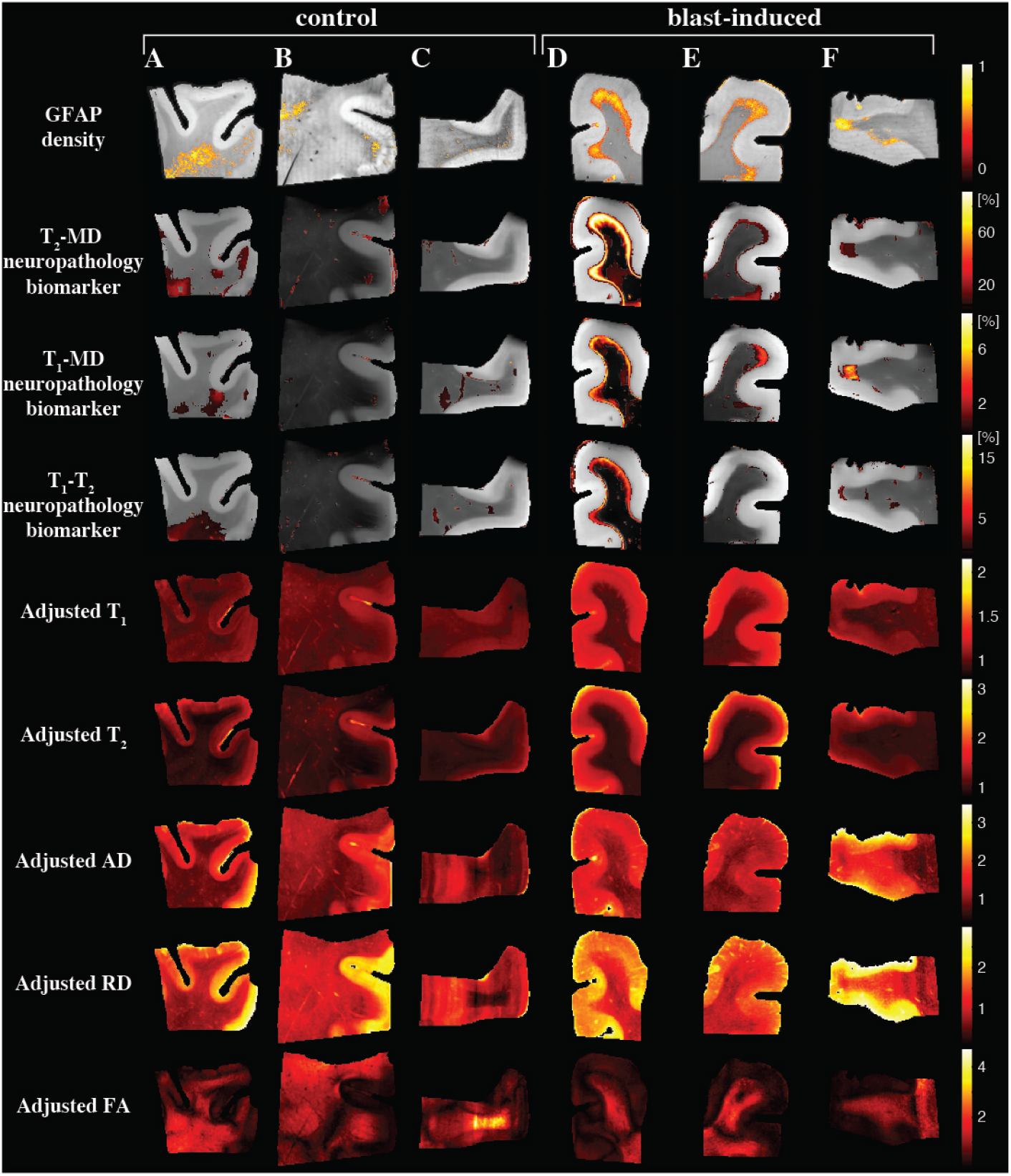
Multidimensional and voxel-averaged MRI maps. (A)-(C) are subjects that were not exposed to blast (Cases 3, 4, and 7), while (D)-(F) were (Cases 10-12). The different rows correspond to the different MRI contrasts, including all the conventional relaxation and DTI parameters, and the proposed multidimensional astrogliosis maps. In addition, the co-registered histological GFAP density maps are shown. Multidimensional neuropathology maps overlaid onto proton density images show substantial injury along the gray-white matter interface, while conventional MRI maps of *T*_1_, *T*_2_, AD, RD, and FA do not show visible abnormalities. Note that to facilitate visualization, the multidimensional neuropathology MRI biomarker maps were thresholded at 10% of the maximal intensity and overlaid on grayscale proton density images.

Qualitatively, the multidimensional MRI neuropathology maps follow closely the GFAP density images, with the *T*_2_-MD maps having a significantly larger dynamic range of intensities as compared with the *T*_1_-MD and *T*_1_-*T*_2_ maps, pointing to increased sensitivity. The standard deviations of the MRI neuropathology biomarkers could be computed from the multidimensional processing framework, thus providing a quantified measure of the biomarker uncertainty. Maps of the standard deviations of the cases in Fig. 5 are shown in Supplementary Figure 4. The standard deviations are under an order of magnitude smaller than the corresponding means, pointing to relative stability and low uncertainty.

Conventional MRI and DTI maps provided useful anatomical macroscopic information, especially gray-white matter separation, however, they fail to capture the microstructural changes that are induced by astrogliosis.

### Strong correlation between MRI measures and GFAP density

The multimodal data set in this study allows one to investigate the strength of the relationships between the multidimensional MRI derived biomarkers we discovered and histologically based astrogliosis and their spatial agreement. A whole image approach (as opposed to ROIs), in which all regions from the MRI maps and histological images were included, was selected to achieve the most objective measures of correlations. After matching the GFAP density images resolution to their MRI counterparts, both MRI and histological maps were downsampled by a factor of 12 to account for co-registration errors and to reduce spatial dependencies (Supplementary Figure 5), resulting in a total of 556 pairs of MR image volumes and GFAP densities from all 14 subjects. Figure 6 summarizes the association between the investigated MR metrics – multidimensional MRI neuropathology biomarker maps and conventional voxel-averaged images – and the pathological findings across the entire images, grouped according to the Case number. This way allows for inferences about both within and between subject correlations (see the Statistical Analysis section for more details).

**Fig. 6.**
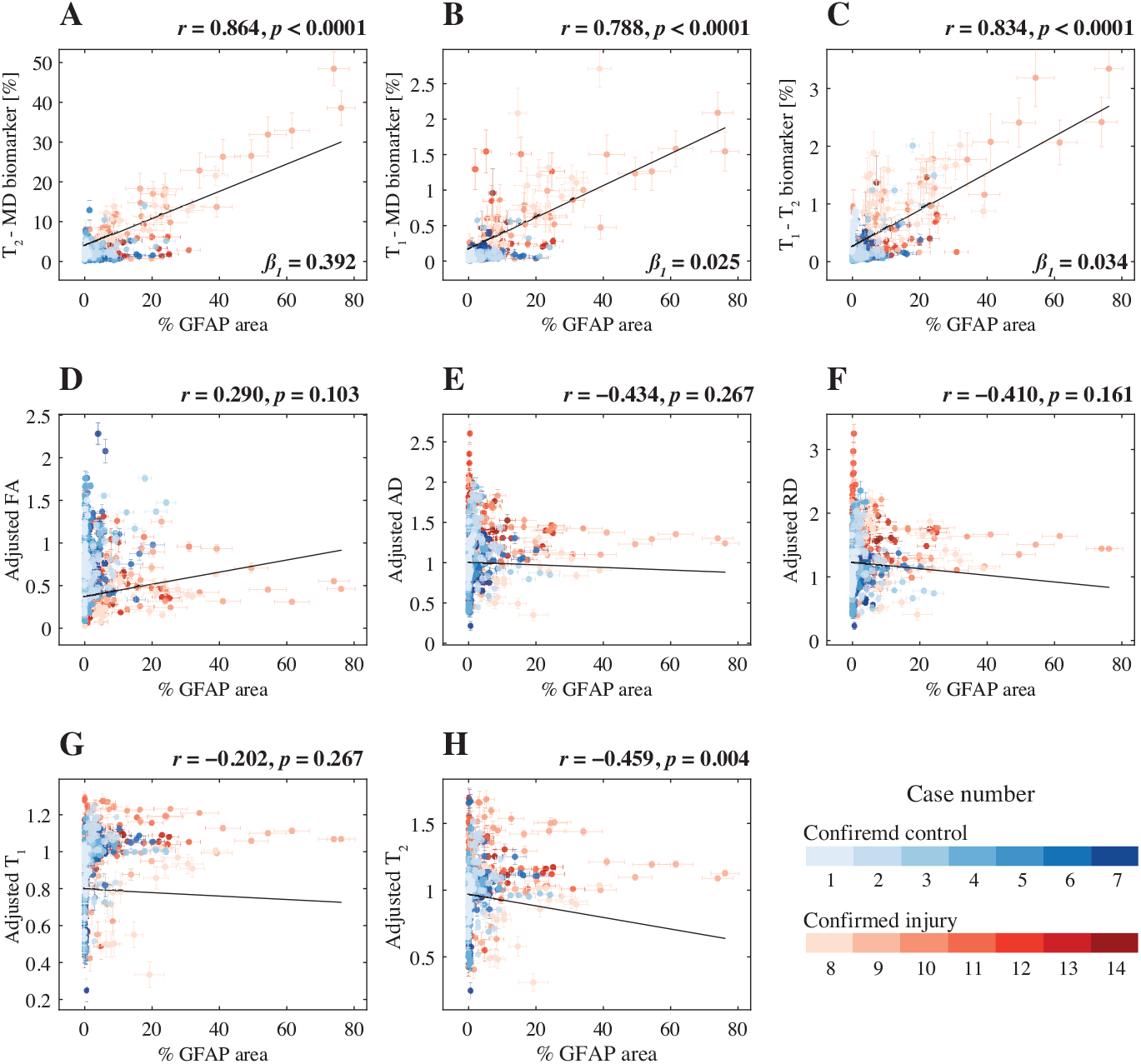
Radiological-pathological correlations between MRI metrics and GFAP density. GFAP density (% area) from 556 tissue regions from 14 subjects (color-coded, see legend) and the corresponding MR parameter correlations. Individual data points represent the mean value from each postmortem tissue sample. Scatterplots of the mean (with 95% confidence interval error bars) % area GFAP and (A) *T*_2_-MD, (B) *T*_1_-MD, and (C) *T*_1_-*T*_2_ injury MRI biomarkers show strong positive and significant correlation with GFAP density. The conventional MRI metrics (D)-(G) did not result in strong and significant correlations with % area GFAP, with the exception of weak yet significant correlation of (H) voxel-averaged *T*_2_.

We found that GFAP density was strongly and significantly correlated with the *T*_2_-MD neuropathology biomarker (*r* = 0.864, *p* < 0.0001), the *T*_1_-MD neuropathology biomarker (*r* = 0.788, *p* < 0.0001), and the *T*_1_-*T*_2_ neuropathology biomarker (*r* = 0.834, *p* < 0.0001). Importantly, these results indicated that higher intensity of the multidimensional neuropathology MRI biomarkers is associated with increased astrogliosis severity, regardless of the tissue type studied. Notably, while the slopes (*β*_1_ in Fig. 6) from the *T*_1_-MD and *T*_1_-*T*_2_ regression analyses indicated potentially low sensitivity (0.025 and 0.034, respectively), the slope from the *T*_2_-MD neuropathology MRI biomarker was an order of magnitude larger (0.392). From the conventional voxel-averaged images, the only measure that had a significant yet weak correlation with GFAP density was the adjusted *T*_2_ (*r* = − 0.459, *p* = 0.004). The effect of the subject’s age was insignificant for all MRI contrasts.

### Astrogliosis is detectable in individuals

We compared multidimensional MRI neuropathology maps intensities from normal-appearing and injured regions to test whether our approach can be used to image astrogliosis in a single subject. Normal-appearing and injured ROIs were defined automatically based on the GFAP density image (e.g., Fig. 4D), and were used as binary masks on the multidimensional MRI neuropathology maps to obtain average and 95% confidence intervals of the intensity values. Figures 7 A-C show these comparisons for the neuropathology MRI biomarkers derived from *T*_2_-MD, *T*_1_-MD, and *T*_1_-*T*_2_, respectively, for all subjects. With the exception of Case 9 for *T*_2_-MD and *T*_1_-MD, and of Case 14 for *T*_1_-*T*_2_, the multidimensional MRI neuropathology maps were shown to be capable of detecting astrogliosis in individuals, illustrated by significant differences between the ROIs (*p* < 0.0001).

**Fig. 7.**
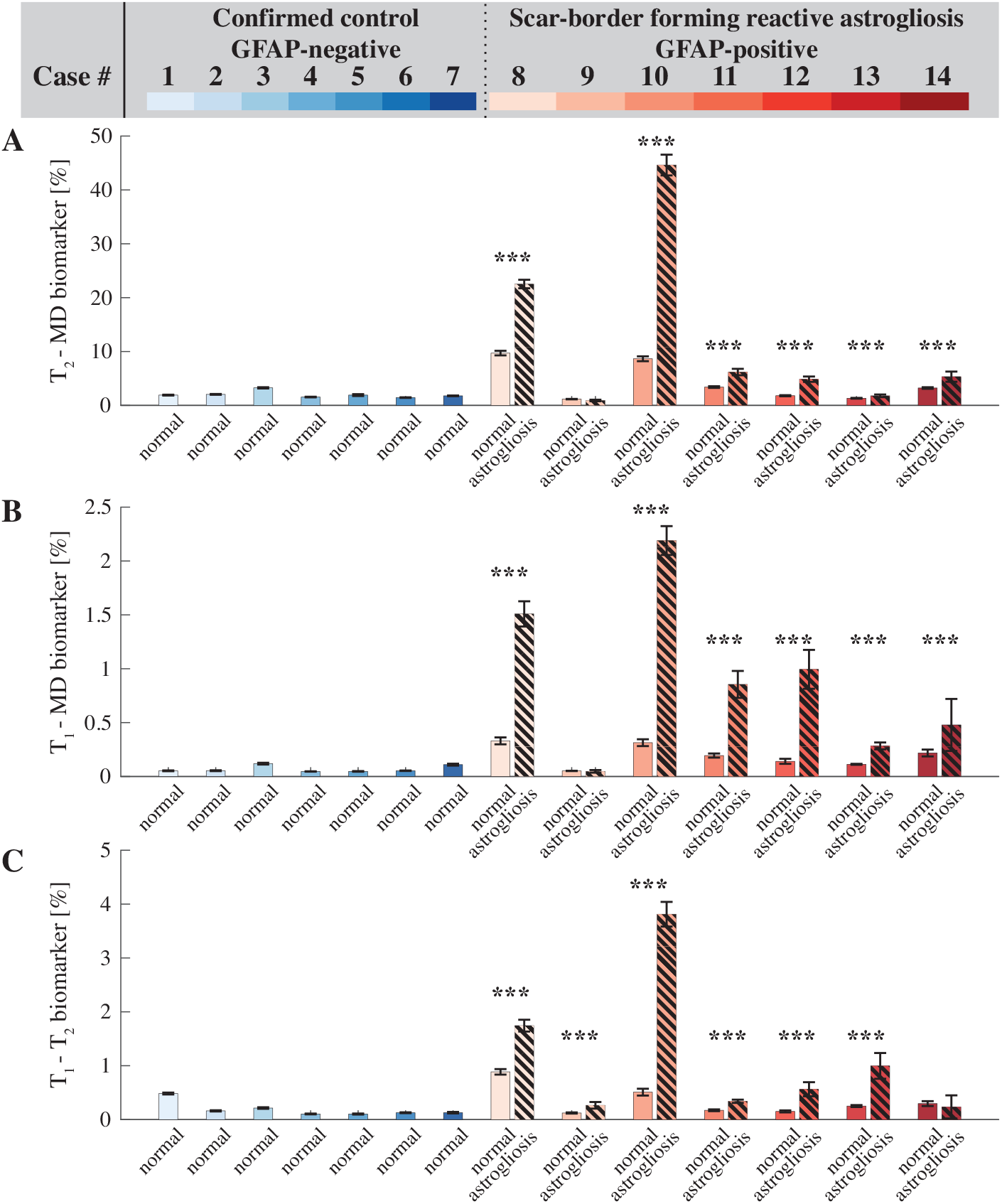
Within-subject comparison of normal-appearing and GFAP-positive regions. Examining the (A) *T*_2_-MD, (B) *T*_1_-MD, and (C) *T*_1_-*T*_2_ neuropathology MRI biomarker maps, areas with increased GFAP density had significantly stronger intensities compared with normative regions.

## Discussion

This is the first report of a noninvasive MRI framework to directly map astrogliosis in individual brains *ex vivo*. In this study we showed that astrogliosis induces microstructural changes that result in a distinct multidimensional MRI spectral signature. Further, we developed a novel approach to utilize this information and obtain MRI maps of astroglial neuropathology in individual human brains. We found that the multidimensional MRI astrogliosis biomarker maps are significantly and strongly correlated with co-registered histological images of increased GFAP expression. We showed that our approach has the spatial sensitivity to detect altered tissue states at the individual level by comparing normal-appearing and histologically confirmed regions within the same brain.

All the injured cases we examined in this study (i.e., chronic blast TBI) had scar-border forming astrogliosis but importantly, absent additional cellular changes, most notably without evidence of tau pathology or axonal injury. This neuropathological uniqueness towards astrogliosis presented an opportunity for a targeted study that provided inferences with a good degree of specificity. Conversely, very few studies have directly investigated the effect astrogliosis has on MRI diffusion and relaxation properties, and those who have did so by using animal injury models that always resulted in substantial axonal damage and other major cellular microstructural changes in addition to gliosis [13–16].

Our results show that scar-border forming astrogliosis was mainly present in superficial WM and involved the gray-white matter junction, in all the cases we examined in this study. Therefore, focusing on changes to the multidimensional spectral signature between normal-appearing WM and regions with astrogliosis could elucidate the chemical and microstructural alterations induced by this type of neuropathology. In terms of *T*_2_ relaxation, severe astrogliosis causes an increase in *T*_2_ compared with normal-appearing WM, and similarly, regions with astrogliosis are characterized by faster MD compared with normal-appearing WM (pink highlight in Figs. 3D and E). Increases in both relaxation times and diffusivities point to a reduction in the degree of axonal packing and density that is expected to result from the presence of highly reactive astrocytes (see Fig. 1) [14, 38]. These measurable microstructural changes can be attributed solely to astrogliosis because no axonal damage or demyelination were histologically observed in any of the cases in the study. In addition, we recently reported that axonal injury has a distinct multidimensional MRI signature characterized by shortening of *T*_1_ and *T*_2_ [31]. These findings along with the current results indicate that decoupling of axonal injury and astrogliosis using a single framework should be possible because of the opposing effects these neuropathologies have on their respective multidimensional MRI spectra (i.e., reduction or increase in *T*_1_ and *T*_2_ for axonal injury and astrogliosis, respectively).

The novel framework we propose here helped to elucidate the underpinning of MRI signal response from astrogliosis, and importantly, showed that no onedimensional *T*_1_, *T*_2_, or diffusion MRI measurement is able to disentangle the microstructural alterations caused by this neuropathology. It is therefore not surprising that very little progress has been made thus far towards the radiological assessment and mapping of astrogliosis, as this study appears to be the first one to use multidimensional MRI to address this problem. In principle, our framework can be readily extended to include more MR dimensions (e.g., diffusion orientation [23] and magnetization transfer [39]) to improve sensitivity, and more histological stains (e.g., axonal damage, myelin) to improve specificity.

DTI metrics are well-studied in the context of microstructural alterations due to TBI [40–42], however, it is becoming increasingly evident that they are inconsistent in their observed response [43–45]. Moreover, one of the only comprehensive MRI studies that examined the contribution of astrogliosis to DTI metrics showed that while significant increase in FA was associated with astrogliosis in cortical GM, significant changes in WM were related to demyelination, and not astrogliosis [14]. These findings are consistent with our results that showed correlations between DTI metrics and astrogliosis in WM were not significant, reflecting the heterogeneity and variability of real life TBI cases, as opposed to animal models.

Although we used histological data to infer the spatial distribution of astrogliosis as an integrated part of our machine learning framework, our approach is not limited to *ex vivo* studies. Using histology to locate normal-appearing regions of the brain can be replaced by collecting baseline multidimensional MRI spectra from healthy participants that will define a normative brain, and then apply our approach to detect abnormalities in the rich data. Acquiring such data in a clinical setting has become feasible following recent developments of multidimensional MRI clinical protocols by multiple groups [28, 46, 47]. As is the case with any other single-patient analysis methods, substantial amounts of normative data will be required to establish a reference ‘atlas’ [48, 49]. Additional limitations and confounds specific to our study include the effects of postmortem decay, fixation and resulting dehydration. The fixation process by itself and the delay in fixation from the time of death (i.e., postmortem interval, PMI) cause changes to tissue properties and affect measured MRI parameters [50], which prevents direct comparison with *in vivo* data. Furthermore, information regarding the PMI was available from only about half of the subjects and cannot be controlled for in our study. Although showing that the GFAP deposition corresponds to changes in the multidimensional signature was only possible using combined *ex vivo* MRI and immunostaining, it remains to be demonstrated *in vivo*.

In summary, being able to selectively focus on sub-voxel relaxation and diffusion components combined with a simple yet effective machine learning approach to detect anomalies in individual subjects provides a framework for mapping of astrogliosis with high precision. While MRI may offer promise to detect subtle microstructural differences at the group level, the goal of clinical neuroimaging is to be applicable at the individual level, potentially facilitating individualized diagnosis and subsequent therapy. This work emphasizes the importance and the potential of combining relaxation and diffusion MRI with artificial intelligence for studying human brain astroglial reactivity noninvasively.

## Methods

### Donor specimens

We evaluated 14 autopsy-derived brain autopsy specimens from two different human brain collections. Formalin-fixed portions of approximately 20 × 20 × 10 × mm^3^ from the frontal lobe were obtained from two civilian subjects enrolled in the Transforming Research and Clinical Knowledge in Traumatic Brain Injury study (TRACK-TBI; https://tracktbi.ucsf.edu/transforming-research-and-clinical-knowledge-tbi) (Cases 2 and 3), and 12 military subjects from the Department of Defense/Uniformed Services University Brain Tissue Repository (DoD/USU BTR, https://www.researchbraininjury.org, Uniformed Services University of the Health Sciences, Bethesda, MD; Cases 1, 4-14). For each case, the next–of-kin or legal representative provided written consent for donation of the brain for use in research. The brain tissues used have undergone procedures for donation of the tissue, its storage, and use of available clinical information that have been approved by the USU Institutional Review Board (IRB) prior to the initiation of the study. All experiments were performed in accordance with current federal, state, DoD, and NIH guidelines and regulations for post-mortem analysis. A detailed description of demographics for the subjects from whom brain tissue samples were obtained is listed in Table 1 and Supplementary Table 1.

Of the total 14 cases evaluated, there were 7 cases with known IAS and 7 control cases negative for IAS, based on prior neuropathologic examination at the DoD/USU BTR. IAS pathology was diagnosed in these cases from microscopic examination of cerebral sections immunostained for GFAP, based on the presence of prominent scar-border forming astrogliosis involving subpial glial plate, penetrating cortical blood vessels, grey-white matter junctions, and structures lining the ventricles, as has been described and published by authors in this study (DPP) [34]. However, initially, tissues from all 14 of these cases were received from the DoD/USU BTR for blinded MRI examination without access to the corresponding histopathology findings, TBI history, or other medical history.

### MRI acquisition

Prior to MRI scanning, each formalin-fixed brain specimen was transferred to a phosphate-buffered saline (PBS) filled container for 12 days to ensure that any residual fixative was removed from the tissue. The specimen was then placed in a 25 mm tube, and immersed in perfluoropolyether (Fomblin LC/8, Solvay Solexis, Italy), a proton free fluid void of a proton-MRI signal. Specimens were imaged using a 7 T Bruker vertical bore MRI scanner equipped with a microimaging probe and a 25 mm quadrupole RF coil.

Multidimensional data were acquired using a 3D inversion recovery diffusion-weighted sequence with a repetition time of 1,000 ms, in-plane resolution of 200*×*200 *μ*m^2^, and slice thickness of 300 *μ*m. To encode the multidimensional MR space spanned by *T*_1_ and *T*_2_ (i.e., *T*_1_-*T*_2_), by *T*_1_ and diffusion (i.e., *T*_1_-MD), and by *T*_2_ and diffusion (i.e., *T*_2_-MD), 56, 302, and 302 images were acquired, respectively, according to a previously published sampling scheme [31, 36]. Additional parameters of the MRI pulse sequence can be found in the Supplementary Material.

A standard DTI imaging protocol was applied with the same imaging parameters as the multidimensional data and using 21 diffusion gradient directions and four b-values ranging from 0 to 1400 s/mm^2^.

Lastly, a high-resolution MRI scan with an isotropic voxel dimension of 100 μm was acquired using a fast low angle shot (FLASH) sequence [51] with a flip angle of 49.6^*◦*^ to serve as a high resolution reference image and facilitate co-registration of histopathological and MR images.

### Histology and Immunohistochemistry

After MRI scanning, each tissue block was transferred for histopathological processing. Tissue blocks from each brain specimen was processed using an automated tissue processor (ASP 6025, Leica Biosystems, Nussloch, Germany). After tissue processing, each tissue block was embedded in paraffin and cut in a series of 5 *μ*m thick consecutive sections. The first section was selected for hematoxylin and eosin (H&E) stains, while the remaining sections were selected for immunohistochemistry procedures using a Leica Bond III automated immunostainer with a diaminobenzidine chromogen detection system (DS9800, Leica Biosystems, Buffalo Grove, IL). Immunohistochemistry was performed for glial fibrillary acidic protein (GFAP) to evaluate presence of astrogliosis, for amyloid precursor protein (APP) for the detection of axonal injury, and for abnormally phosphorylated tau (AT8) protein. Two sections per antibody were stained at 300 μm apart from each other, in accordance with the MRI slice thickness. More details regarding immunohistochemistry can be found in the Supplementary Information.

All stained sections were digitally scanned using an Aperio scanner system (Aperio AT2 - High Volume, Digital whole slide scanning scanner, Leica Biosystems, Inc., Richmond, IL) and stored in Biolucida, a hub for 2D and 3D image data (MBF Bioscience, Williston, VT) for further assessment and analyses. A Zeiss Imager A2 (ImagerA2 microscope, Zeiss, Munich, Germany) bright-field microscope with × 40 and × 63 magnification lenses was used to identify and photograph histologic and pathologic details, as needed.

### Quantification of astrogliosis

Images of GFAP-stained sections were digitized using an Aperio whole slide scanning scanner system (Leica Biosystems, Richmond, IL) at × 20 magnification. The following steps, all implemented using MATLAB (The Mathworks, Natick, MA), were taken to allow for a quantitative analysis of the GFAP images. First, the images were transformed into a common, normalized space to enable improved quantitative analysis [52]. Then, the normalized images were deconvolved to unmix the primary (GFAP) and secondary (H&E) stains, and background to three separate channels [53]. Once an GFAP-only image has been obtained, a final thresholding step individualized for each slice was taken to exclude non-specific staining and to allow for a subsequent % area calculation.

### Histology-MRI co-registration

The high-resolution MR images were used as anatomical references to which the histological images were registered to. Areas in the histological images that grossly diverged from the wet tissue state (i.e., the MR images) due to deformation were manually removed, while maintaining the image aspect ratio. Following convergence of 2D affine co-registration of histology and MR images (Image Processing Toolbox, MATLAB, The Mathworks, Natick, MA), we performed a 2D diffeomorphic registration refinement between the GFAP image slices and MRI volumes. This was done to recover true in-plane tissue shape and bridge over residual differences between the modalities. The diffeomorphic registration procedure in this study was performed using an efficient implementation of the greedy diffeomorphic algorithm [54], provided as an open-source software package (*greedy*, https://github.com/pyushkevich/greedy). The greedy software was initialized and used as previously described [55]. The transformed histology images were overlaid on MR images to assess the quality of the coregistration, and the Jaccard index [56] was computed to quantify the overlap scores between the co-registered modalities (Supplementary Figure 6).

### *T*_1_ and *T*_2_ maps and diffusion tensor MRI processing

Diffusion tensor imaging parameters [57], axial diffusivity (AD), radial diffusivity (RD), and fractional anisotropy (FA), were calculated using in-house MATLAB (The Mathworks, Natick, MA) code based on previous work [58].

Conventional quantitative relaxation maps were first computed by fitting the signal decay to monoexponential functions. The *T*_1_ value was computed by fitting a subset of the multidimensional data that included 20 images with inversion times in the range of 12 ms and 980 ms. The *T*_2_ value was computed by fitting a subset of the multidimensional data that included 20 images with echo times in the range of 10.5 ms and 125 ms.

We also applied a commonly used strategy [41, 59] to correct for possible between-subject differences arising from postmortem effects; we adjusted each voxel-averaged MRI parameter by dividing them by the mean for that parameter across all the normal-appearing WM voxels in each brain sample.

### Image domain masks

The FA maps were used to derive WM and GM image masks (defined using a threshold of 0.2). The co-registered GFAP density image and its inverse image (i.e., complementary binary image) were used as ‘injured’ and ‘normal’ image domain masks, respectively. Normal-appearing WM and GM image domain masks were obtained by multiplying the WM and GM masks with the ‘normal’ tissue mask. All image masks were eroded using a disk-shaped structuring element with a radius of 1 to avoid partial volume bias from adjacent structures or from the edges of the brain tissue block.

### Multidimensional MRI processing

Prior to processing, multidimensional MRI data were denoised using an adaptive nonlocal multispectral filter (i.e., NESMA [60]), which was shown to reduce noise and improve the accuracy of the resulting injury MRI biomarker maps [61]

The filtered data were then processed using a marginally-constrained, *ℓ*_2_-regularized, nonnegative least square optimization to compute the multidimensional distribution in each voxel, as previously described [27, 31, 36]. It is a well-tested approach that had been proved robust and reliable [62–65], which in this study had resulted in three types of distributions in each voxel: *T*_1_-*T*_2_, *T*_2_-MD, and *T*_1_-MD.

We implemented the following procedure to correct for possible between-subject differences arising from postmortem effects: First, the normal-appearing WM mask was applied, and the maximal peak location in the spectral domain (e.g., *T*_1_-*T*_2_) was automatically found (this step was repeated for each subject). A control subject was then selected to be used as a reference (Case 1 in this study), to which all the remaining cases were aligned to in the spectral domain. This procedure ensures standardization across subjects, equivalent to the well-established strategy employed for voxel-average images, in which image values are divided by the mean across all the normal-appearing WM voxels in each brain sample [41, 59].

### Statistical analysis

Linear mixed-effects model framework was used to study correlations between MRI-derived maps and GFAP density images. Random effects were added to model the within-subject correlation among histological samples. A whole image approach (as opposed to ROIs), in which all regions from the MRI maps and histological images were included, was used to achieve the most objective measures of correlations. Potential spatial correlation and co-dependencies within subjects were accounted for in two ways: (1) both MRI and histological maps were downsampled by a factor of 12 to reduce spatial dependencies [66], and (2) a Gaussian structured noise covariance matrix was included in the linear mixed-effects regression process to account for the spatial correlations among voxels. Subject age was included as covariate. A two-sample t-test was performed on all pairs of ROIs (i.e., normal-appearing and astrogliosis) to determine whether they are significantly different from one another.

False discovery rate (FDR) correction was carried out to take the overall number of pairwise contrasts into account [67]. A P-value of 0.05 was considered statistically significant. R was used for the computation.

## Data availability

The datasets generated and analyzed during the current study are available from the corresponding author on reasonable request.

### Supplementary information

Supplementary information is available online.

## Acknowledgments

We thank the subjects’ families that consented for brain donations for the better understanding of TBI consequences. The authors thank Dr. Murat Bilgel for insights into the statistical analysis. We also thank Patricia Lee, Nichelle Gray and Paul Gegbeh for their valuable technical work. We are grateful to Stacey Gentile, Deona Cooper and Harold Kramer Anderson for their administrative assistance. We thank the TRACK-TBI Investigators. This research was partially supported by a grant from the U.S. Department of Defense, Program Project 308430 USUHS. Support for this work also included funding from the U.S. Department of Defense to the Brain Tissue Repository and Neuropathology Core, Center for Neuroscience and Regenerative Medicine (CNRM). DB was supported by the CNRM Neuroradiology-Neuropathology Correlation Core. DP, DPP, and DLB were supported by the CNRM and USUHS. DB was supported by the Intramural Research Program of the National Institute on Aging. PJB was supported by the Intramural Research Program of the *Eunice Kennedy Shrive*r National Institute of Child Health and Human Development.

## Declarations

The opinions expressed herein are those of the authors and are not necessarily representative of those of the Uniformed Services University of the Health Sciences (USUSH), the Department of Defense (DOD), the NIH or any other US government agency.

## References

[1] Sofroniew, M. V. & Vinters, H. V. Astrocytes: biology and pathology. Acta Neuropathologica 119, 7–35 (2010). https://doi.org/10.1007/s00401-009-0619-8.

[2] Bush, T. G. et al. Leukocyte infiltration, neuronal degeneration, and neurite outgrowth after ablation of scar-forming, reactive astrocytes in adult transgenic mice. Neuron 23, 297–308 (1999). https://doi.org/10.1016/S0896-6273(00)80781-3.

[3] Wanner, I. B. et al. Glial scar borders are formed by newly proliferated, elongated astrocytes that interact to corral inflammatory and fibrotic cells via stat3-dependent mechanisms after spinal cord injury. Journal of Neuroscience 33, 12870–12886 (2013). https://doi.org/10.1523/JNEUROSCI.2121-13.2013.

[4] Voskuhl, R. R. et al. Reactive astrocytes form scar-like perivascular barriers to leukocytes during adaptive immune inflammation of the cns. Journal of Neuroscience 29, 11511–11522 (2009). https://doi.org/10.1523/JNEUROSCI.1514-09.2009.

[5] Pekny, M. & Pekna, M. Astrocyte reactivity and reactive astrogliosis: Costs and benefits. Physiological Reviews 94, 1077–1098 (2014). https://doi.org/10.1152/physrev.00041.2013.

[6] Heiland, D. H. et al. Tumor-associated reactive astrocytes aid the evolution of immunosuppressive environment in glioblastoma. Nature Communications 10, 2541 (2019). https://doi.org/10.1038/s41467-019-10493-6.

[7] Shao, W. et al. Suppression of neuroinflammation by astrocytic dopamine d2 receptors via b-crystallin. Nature 494, 90–94 (2013). https://doi.org/10.1038/nature11748.

[8] Sofroniew, M. V. Astrocyte barriers to neurotoxic inflammation. Nature Reviews Neuroscience 16, 249–263 (2015). https://doi.org/10.1038/nrn3898.

[9] Escartin, C., Guillemaud, O. & Sauvage, M. C. Questions and (some) answers on reactive astrocytes. Glia 67, 2221–2247 (2019). https://doi.org/10.1002/glia.23687.

[10] Anderson, M. A. et al. Astrocyte scar formation aids central nervous system axon regeneration. Nature 532, 195–200 (2016). https://doi.org/10.1038/nature17623.

[11] Herrmann, J. E. et al. Stat3 is a critical regulator of astrogliosis and scar formation after spinal cord injury. Journal of Neuroscience 28, 7231–7243 (2008). https://doi.org/10.1523/JNEUROSCI.1709-08.2008.

[12] Wheeler, M. A. et al. Mafg-driven astrocytes promote cns inflammation. Nature 578, 593–599 (2020). https://doi.org/10.1038/s41586-020-1999-0.

[13] Schwartz, E. D., Duda, J., Shumsky, J. S., Cooper, E. T. & Gee, J. Spinal cord diffusion tensor imaging and fiber tracking can identify white matter tract disruption and glial scar orientation following lateral funiculotomy. Journal of Neurotrauma 22, 1388–1398 (2005). https://doi.org/10.1089/neu.2005.22.1388.

[14] Budde, M. D., Janes, L., Gold, E., Turtzo, L. C. & Frank, J. A. The contribution of gliosis to diffusion tensor anisotropy and tractography following traumatic brain injury: validation in the rat using fourier analysis of stained tissue sections. Brain 134, 2248–2260 (2011). https://doi.org/10.1093/brain/awr161.

[15] Zhuo, J. et al. Diffusion kurtosis as an in vivo imaging marker for reactive astrogliosis in traumatic brain injury. NeuroImage 59, 467–477 (2012). https://doi.org/10.1016/j.neuroimage.2011.07.050.

[16] Chary, K. et al. Quantitative susceptibility mapping of the rat brain after traumatic brain injury. NMR in Biomedicine 34(2021). https://doi.org/10.1002/nbm.4438.

[17] Ishiki, A. et al. Neuroimaging-pathological correlations of [18f]thk5351 pet in progressive supranuclear palsy. Acta Neuropathologica Communications 6, 53 (2018). https://doi.org/10.1186/s40478-018-0556-7.

[18] Hatakeyama, T. et al. Temporal and spatial changes in reactive astrogliosis examined by 18f-thk5351 positron emission tomography in a patient with severe traumatic brain injury. European Journal of Hybrid Imaging 5, 26 (2021). https://doi.org/10.1186/s41824-021-00121-2.

[19] Benjamini, D. & Basser, P. J. Multidimensional correlation mri. NMR in Biomedicine 33(2020). https://doi.org/10.1002/nbm.4226.

[20] Slator, P. J. et al. Combined diffusion-relaxometry microstructure imaging: Current status and future prospects. Magnetic Resonance in Medicine 86, 2987–3011 (2021). https://doi.org/10.1002/mrm.28963.

[21] Benjamini, D. & Basser, P. Use of marginal distributions constrained optimization (madco) for accelerated 2d mri relaxometry and diffusometry. Journal of Magnetic Resonance 271, 40–45 (2016). https://doi.org/10.1016/j.jmr.2016.08.004.

[22] Kim, D., Doyle, E. K., Wisnowski, J. L., Kim, J. H. & Haldar, J. P. Diffusion-relaxation correlation spectroscopic imaging: A multidimensional approach for probing microstructure. Magnetic Resonance in Medicine 78, 2236–2249 (2017). https://doi.org/10.1002/mrm.26629.

[23] Topgaard, D. Multidimensional diffusion mri. Journal of Magnetic Resonance 275, 98–113 (2017). https://doi.org/10.1016/j.jmr.2016.12.007.

[24] Slator, P. J. et al. Data-driven multi-contrast spectral microstructure imaging with inspect: Integrated spectral component estimation and mapping. Medical Image Analysis 71, 102045 (2021). https://doi.org/10.1016/j.media.2021.102045.

[25] Hutter, J. et al. Integrated and efficient diffusion-relaxometry using zebra. Scientific Reports 8, 15138 (2018). https://doi.org/10.1038/s41598-018-33463-2.

[26] Manhard, M. K. et al. A multi-inversion multi-echo spin and gradient echo echo planar imaging sequence with low image distortion for rapid quantitative parameter mapping and synthetic image contrasts. Magnetic Resonance in Medicine 86, 866–880 (2021). https://doi.org/10.1002/mrm.28761.

[27] Benjamini, D. & Basser, P. J. Magnetic resonance microdynamic imaging reveals distinct tissue microenvironments. NeuroImage 163, 183–196 (2017). https://doi.org/10.1016/j.neuroimage.2017.09.033.

[28] Slator, P. J. et al. Combined diffusion-relaxometry mri to identify dysfunction in the human placenta. Magnetic Resonance in Medicine 82, 95–106 (2019). https://doi.org/10.1002/mrm.27733.

[29] de Almeida Martins, J. P. et al. Computing and visualising intravoxel orientation-specific relaxation–diffusion features in the human brain. Human Brain Mapping 42, 310–328 (2021). https://doi.org/10.1002/hbm.25224.

[30] Reymbaut, A. et al. Toward nonparametric diffusion-characterization of crossing fibers in the human brain. Magnetic Resonance in Medicine 85, 2815–2827 (2021). https://doi.org/10.1002/mrm.28604.

[31] Benjamini, D. et al. Diffuse axonal injury has a characteristic multidimensional mri signature in the human brain. Brain 144, 800–816 (2021). https://doi.org/10.1093/brain/awaa447.

[32] Johnson, V. E., Stewart, W. & Smith, D. H. Axonal pathology in traumatic brain injury. Experimental Neurology 246, 35–43 (2013). https://doi.org/10.1016/j.expneurol.2012.01.013.

[33] Phipps, H. et al. Characteristics and impact of u.s. military blast-related mild traumatic brain injury: A systematic review. Frontiers in Neurology 11(2020). https://doi.org/10.3389/fneur.2020.559318.

[34] Shively, S. B. et al. Characterisation of interface astroglial scarring in the human brain after blast exposure: a post-mortem case series. The Lancet Neurology 15, 944–953 (2016). https://doi.org/10.1016/S1474-4422(16)30057-6.

[35] Schwerin, S. C. et al. Expression of gfap and tau following blast exposure in the cerebral cortex of ferrets. Journal of Neuropathology Experimental Neurology 80, 112–128 (2021). https://doi.org/10.1093/jnen/nlaa157.

[36] Pas, K., Komlosh, M. E., Perl, D. P., Basser, P. J. & Benjamini, D. Retaining information from multidimensional correlation mri using a spectral regions of interest generator. Scientific Reports 10, 3246 (2020). https://doi.org/10.1038/s41598-020-60092-5.

[37] Xu, Q.-S. & Liang, Y.-Z. Monte carlo cross validation. Chemometrics and Intelligent Laboratory Systems 56, 1–11 (2001). https://doi.org/10.1016/S0169-7439(00)00122-2.

[38] Mac Donald, C. L. et al. Longitudinal neuroimaging following combat concussion: sub-acute, 1 year and 5 years post-injury. Brain Communications 1(2019). https://doi.org/10.1093/braincomms/fcz031.

[39] Sled, J. G. & Pike, G. B. Quantitative imaging of magnetization transfer exchange and relaxation properties in vivo using mri. Magnetic Resonance in Medicine 46, 923–931 (2001). https://doi.org/10.1002/mrm.1278.

[40] Mac Donald, C. L., Dikranian, K., Bayly, P., Holtzman, D. & Brody, D. Diffusion tensor imaging reliably detects experimental traumatic axonal injury and indicates approximate time of injury. Journal of Neuroscience 27, 11869–11876 (2007). https://doi.org/10.1523/JNEUROSCI.3647-07.2007.

[41] Holleran, L. et al. Axonal disruption in white matter underlying cortical sulcus tau pathology in chronic traumatic encephalopathy. Acta Neuropathologica 133, 367–380 (2017). https://doi.org/10.1007/s00401-017-1686-x.

[42] Bourke, N. J. et al. Traumatic brain injury: a comparison of diffusion and volumetric magnetic resonance imaging measures. Brain Communications 3(2021). https://doi.org/10.1093/braincomms/fcab006.

[43] Budde, M. D. & Annese, J. Quantification of anisotropy and fiber orientation in human brain histological sections. Frontiers in Integrative Neuroscience 7(2013). https://doi.org/10.3389/fnint.2013.00003.

[44] Kamnaksh, A. et al. Diffusion tensor imaging reveals acute subcortical changes after mild blast-induced traumatic brain injury. Scientific Reports 4, 4809 (2015). https://doi.org/10.1038/srep04809.

[45] Dennis, E. L. et al. Enigma military brain injury: A coordinated metaanalysis of diffusion mri from multiple cohorts, 1386–1389 (IEEE, 2018).

[46] Avram, A. V., Sarlls, J. E. & Basser, P. J. Whole-brain imaging of subvoxel t1-diffusion correlation spectra in human subjects. Frontiers in Neuroscience 15(2021). https://doi.org/10.3389/fnins.2021.671465.

[47] Martin, J. et al. Nonparametric d-r1-r2 distribution mri of the living human brain. NeuroImage 245, 118753 (2021). https://doi.org/10.1016/j.neuroimage.2021.118753.

[48] Chamberland, M. et al. Detecting microstructural deviations in individuals with deep diffusion mri tractometry. Nature Computational Science 1, 598–606 (2021). https://doi.org/10.1038/s43588-021-00126-8.

[49] Jolly, A. E. et al. Detecting axonal injury in individual patients after traumatic brain injury. Brain 144, 92–113 (2021). https://doi.org/10.1093/brain/awaa372.

[50] Shepherd, T. M. et al. Postmortem interval alters the water relaxation and diffusion properties of rat nervous tissue — implications for mri studies of human autopsy samples. NeuroImage 44, 820–826 (2009). https://doi.org/10.1016/j.neuroimage.2008.09.054.

[51] Matthaei, D., Frahm, J., Haase, A. & Hanicke, W. Regional physiological functions depicted by sequences of rapid magnetic resonance images. The Lancet 326, 893 (1985). https://doi.org/10.1016/S0140-6736(85)90158-8.

[52] Macenko, M. et al. A method for normalizing histology slides for quantitative analysis, 1107–1110 (IEEE, 2009).

[53] Ruifrok, A. C. & Johnston, D. A. Quantification of histochemical staining by color deconvolution. Analytical and Quantitative Cytology and Histology 23, 291—299 (2001).

[54] Joshi, S., Davis, B., Jomier, M. & Gerig, G. Unbiased diffeomorphic atlas construction for computational anatomy. NeuroImage 23, S151–S160 (2004). https://doi.org/10.1016/j.neuroimage.2004.07.068.

[55] Adler, D. H. et al. Characterizing the human hippocampus in aging and alzheimer’s disease using a computational atlas derived from ex vivo mri and histology. Proceedings of the National Academy of Sciences 115, 4252–4257 (2018). https://doi.org/10.1073/pnas.1801093115.

[56] Jaccard, P. The distribution of the flora in the alpine zone. New Phytologist 11, 37–50 (1912). https://doi.org/10.1111/j.1469-8137.1912.tb05611.x.

[57] Basser, P. J., Mattiello, J. & LeBihan, D. Mr diffusion tensor spectroscopy and imaging. Biophysical Journal 66, 259–67 (1994). https://doi.org/10.1016/S0006-3495(94)80775-1.

[58] Barmpoutis, A. & Vemuri, B. C. A unified framework for estimating diffusion tensors of any order with symmetric positive-definite constraints, 1385–1388 (IEEE, 2010).

[59] Cooper, G. et al. Standardization of t1w/t2w ratio improves detection of tissue damage in multiple sclerosis. Frontiers in Neurology 10(2019). https://doi.org/10.3389/fneur.2019.00334.

[60] Bouhrara, M., Bonny, J.-M., Ashinsky, B. G., Maring, M. C. & Spencer, R. G. Noise estimation and reduction in magnetic resonance imaging using a new multispectral nonlocal maximum-likelihood filter. IEEE Transactions on Medical Imaging 36, 181–193 (2017). https://doi.org/10.1109/TMI.2016.2601243.

[61] Benjamini, D. et al. Multidimensional mri for characterization of subtle axonal injury accelerated using an adaptive nonlocal multispectral filter. Frontiers in Physics 9(2021). https://doi.org/10.3389/fphy.2021.737374.

[62] Provencher, S. W. A constrained regularization method for inverting data represented by linear algebraic or integral equations. Computer Physics Communications 27, 213–227 (1982). https://doi.org/10.1016/0010-4655(82)90173-4.

[63] Kroeker, R. M. & Henkelman, M. R. Analysis of biological nmr relaxation data with continuous distributions of relaxation times. Journal of Magnetic Resonance (1969) 69, 218–235 (1986). https://doi.org/10.1016/0022-2364(86)90074-0.

[64] Mitchell, J., Chandrasekera, T. C. & Gladden, L. F. Numerical estimation of relaxation and diffusion distributions in two dimensions. Progress in Nuclear Magnetic Resonance Spectroscopy 64, 34–50 (2012). https://doi.org/10.1016/j.pnmrs.2011.07.002.

[65] Celik, H., Bouhrara, M., Reiter, D. A., Fishbein, K. W. & Spencer, R. G. Stabilization of the inverse laplace transform of multiexponential decay through introduction of a second dimension. Journal of Magnetic Resonance 236, 134–139 (2013). https://doi.org/10.1016/j.jmr.2013.07.008.

[66] Hyun, D., Crowley, A. L. C. & Dahl, J. J. Efficient strategies for estimating the spatial coherence of backscatter. IEEE Transactions on Ultrasonics, Ferroelectrics, and Frequency Control 64, 500–513 (2017). https://doi.org/10.1109/TUFFC.2016.2634004.

[67] Benjamini, Y. & Yekutieli, D. The control of the false discovery rate in multiple testing under dependency. The Annals of Statistics 29(2001). https://doi.org/10.1214/aos/1013699998.

