## Supplementary Information for "Astrogliosis mapping in individual brains using multidimensional MRI"

### Supplementary Information: Astrogliosis mapping in individual brains using multidimensional MRI

### Supplementary Table and Figures

**Table S1** Medical and neuropsychiatric history in subjects with history of traumatic brain injury (TBI) and healthy controls

| Case | Medical history | Neuropsychiatric history |
|---|---|---|
| 1 | Hyperlipidemia, sleep apnea, shingles, retinal tear | None reported |
| 2 | Alcohol abuse, diabetes | Depression |
| 3 | None reported | None reported |
| 4 | Hypertension, congestive heart failure, aortic valve replacement,diabetes, COPD | None reported |
| 5 | Sports injuries: torn retina, torn patellar tendon, torn ACL | None reported |
| 6 | Alcohol abuse | None reported |
| 7 | Diabetes, hypertension, hypercholesterolemia | None reported |
| 8 | Soft tissue injury, chronic pain | PTSD, reduced cognitive function, confusion, memory loss, seizures, paranoia, delusions, emotional liability |
| 9 | Soft tissue injury, chronic pain, low testosterone, alcohol abuse | PTSD, anxiety, depression |
| 10 | Alcohol and polysubstance abuse, chronic pain, urinary tract infection | PTSD, anxiety, depression |
| 11 | Cannabis abuse, alcohol abuse, low testosterone | PTSD, unspecified bipolar/paranoid disorder |
| 12 | None reported | Sleep disturbance, anxiety, depression |
| 13 | Hypertension, hyperlipidemia, coronary artery disease | None reported |
| 14 | None reported | PTSD, anxiety, depression |

Chronic obstructive pulmonary disease (COPD), Anterior cruciate ligament (ACL), Post traumatic stress disorder (PTSD)

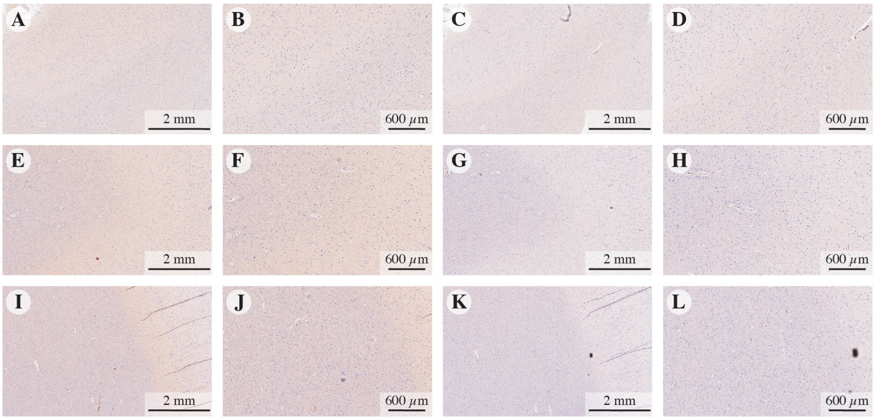


**Fig. S1** Amyloid precursor protein (APP) and phosphorylated tau (pTau) immunoreactivity in representative cases. Subject with impact TBI but without blast exposure (Case 3) (A)-(B) APP, (C)-(D) pTau. Subject without impact TBI but with blast exposure (Case 11) (E)-(F) APP, (G)-(H) pTau. Subject with both impact and blast exposure TBI (Case 8) (I)-(J) APP, (K)-(L) pTau. No axonal damage (APP) or tau pathology (pTau) were observed in any othe the cases in this study.

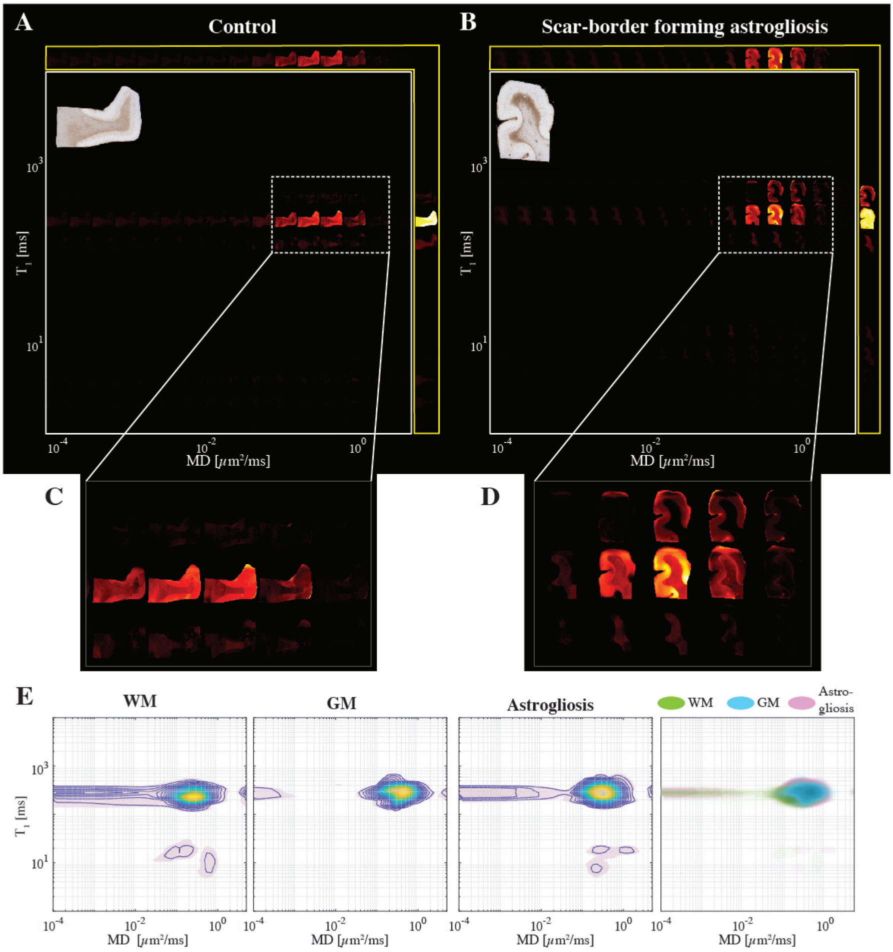


**Fig. S2** Changes in the $T_1$-MD multidimensional MR signature induced by confirmed astrogliosis. Maps of 2D spectra of subvoxel $T_1$-MD values reconstructed on a 16×16 grid of a representative (A) control (case 7) and (B) injured (case 10) subjects, along with their respective GFAP histological image (top left of each panel). Magnified spectral regions showing highest content of spectral information from (C) control and (D) injured cases. (E) $T_1$-MD spectra averaged across all subjects in WM, GM, and GFAP-positive regions of interest (ROIs, left to right), and a superposition of the average spectra from the three ROIs.

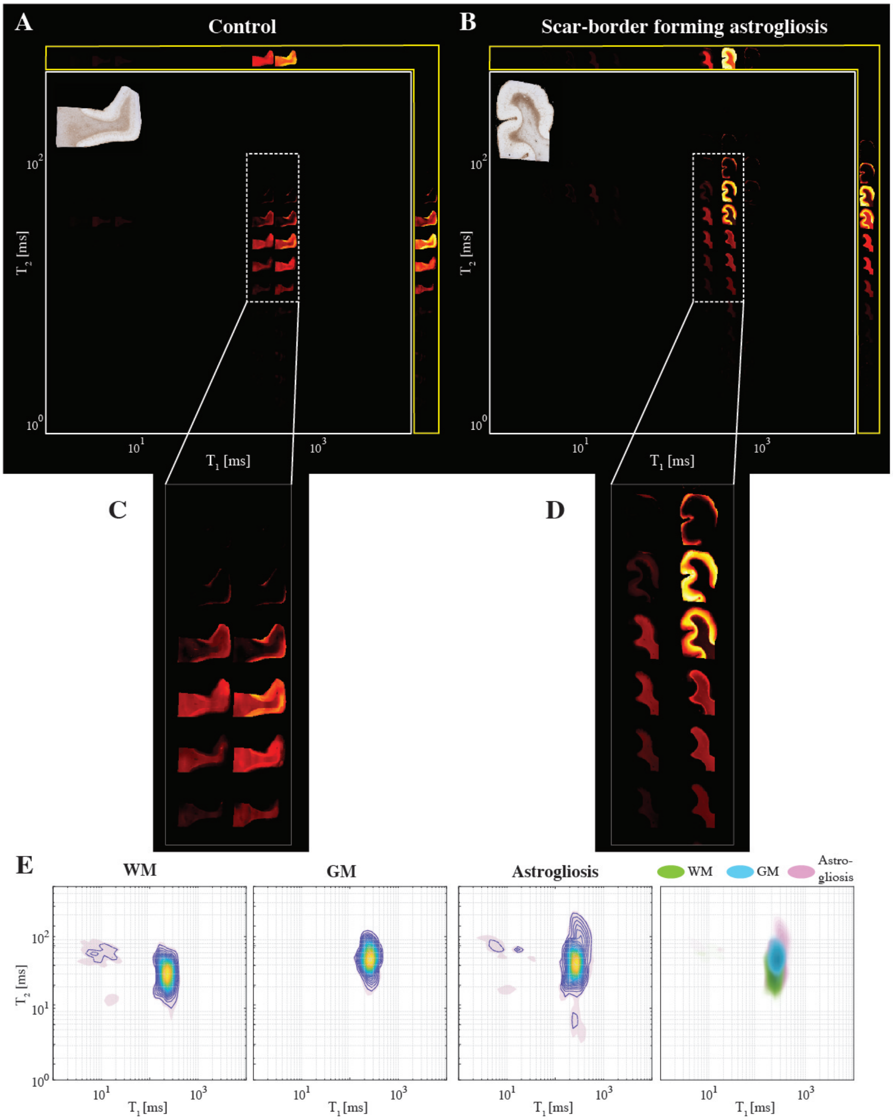


**Fig. S3** Changes in the $T_1$-$T_2$ multidimensional MR signature induced by confirmed astrogliosis. Maps of 2D spectra of subvoxel $T_1$-$T_2$ values reconstructed on a 16×16 grid of a representative (A) control (case 7) and (B) injured (case 10) subjects, along with their respective GFAP histological image (top left of each panel). Magnified spectral regions showing highest content of spectral information from (C) control and (D) injured cases. (E) $T_1$-$T_2$ spectra averaged across all subjects in WM, GM, and GFAP-positive regions of interest (ROIs, left to right), and a superposition of the average spectra from the three ROIs.

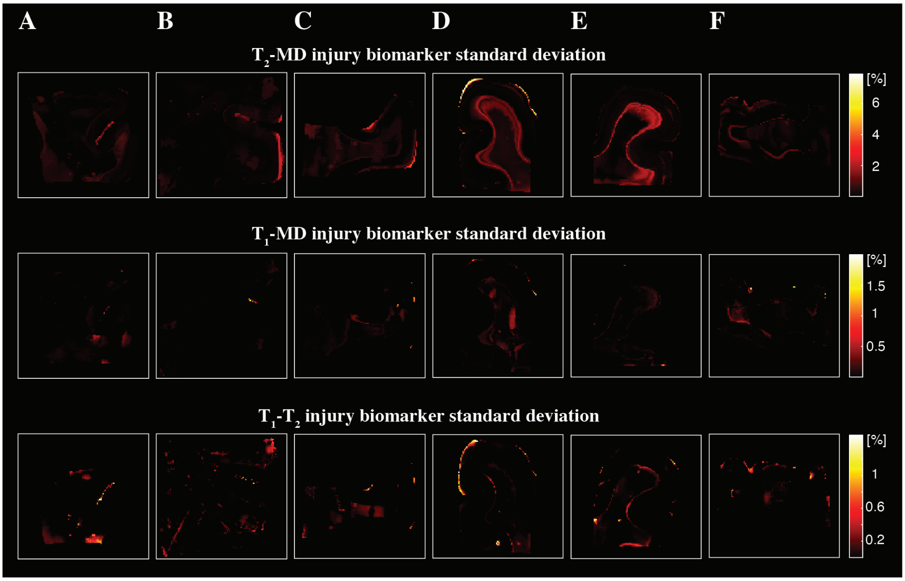


**Fig. S4** Maps of injury biomarkers standard deviations from the cases in Fig. 5. The standard deviations were computed from the Monte Carlo cross-validation procedure for anomaly detection in individuals. (A)-(C) are subjects that were not exposed to blast (Cases 3, 4, and 7), while (D)-(F) were (Cases 10-12). The standard deviations are under an order of magnitude smaller than the corresponding means, pointing to relative stability and low uncertainty.

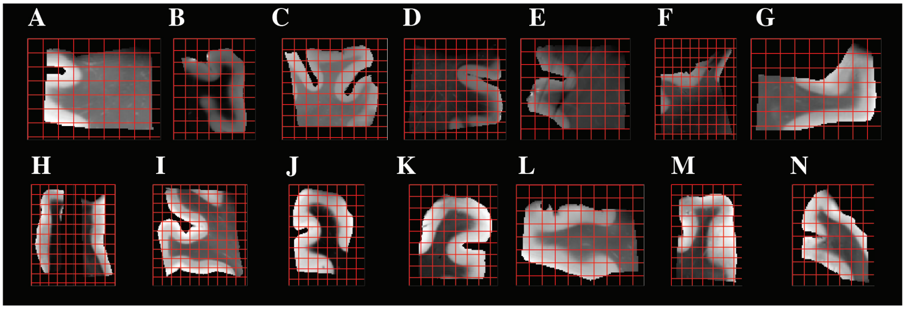


**Fig. S5** Downsampled MR images from all cases, used for whole image correlation. After matching the GFAP density images resolution to their MRI counterparts, both MRI and histological maps were downsampled by a factor of 12 to account for co-registration errors and to reduce spatial dependencies, resulting in a total of 556 pairs of MR image volumes and GFAP densities from all 14 subjects.

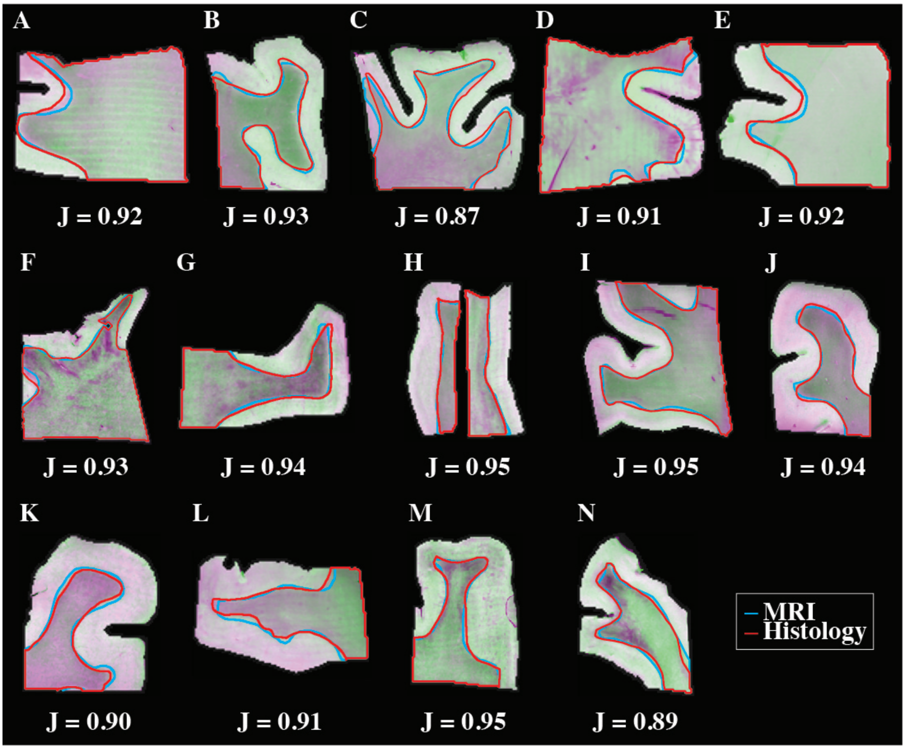


**Fig. S6** MRI and histopathology co-registration qualitative and quantitative accuracy assessments. The transformed histology images were overlaid on MR images using independent intensity scales (purple for GFAP and green for MRI) to assess the quality of the co-registration. The Jaccard index, $J$, was computed to quantify the overlap scores between the co-registered modalities. In addition, qualitative agreement is demonstrated by the overlap in WM-GM interfaces of the MRI and GFAP histology images, which are shown in blue and red, respectively.

### Supplementary Methods

#### MRI acquisition

The acquisition of multidimensional data was done using echo planar imaging (EPI) readout according to the MADCO framework encoding scheme [1, 2], and by varying the following three experimental parameters: the inversion time, $\tau_1$, the echo time, $\tau_2$, and the diffusion weighting, $b$, providing $T_1$-, $T_2$-, and diffusion-weighting, respectively.

The minimal values of the two timing parameters, $\tau_1$ and $\tau_2$, depend on the sample physical dimensions because of the varying imaging matrix size that is intended to keep the spatial resolution constant at 200 $\mu$m in-plane and 300 $\mu$m slice thickness. We kept the minimal $\tau_1$ and $\tau_2$ relatively constant across the samples at $\tau_1 = 12.0 \pm 1.5$ and $\tau_2 = 10.5 \pm 0.8$, by adjusting the number of EPI segments as necessary (<8).

The three 1D distributions of $T_1$, $T_2$, and MD, were estimated, respectively, with the following data acquisition protocols: A 1D $T_1$-weighted data set ($b = 0$, $\tau_2 = 10.5$ ms) with 20 logarithmically sampled $\tau_1$ values ranging from 12 to 980 ms by using an IR-DWI-EPI sequence; a 1D $T_2$-weighted data set ($b$=0) with 20 logarithmically sampled $\tau_2$ values ranging from 10.5 to 125 ms by using a DWI-EPI sequence. For diffusion encoding, we used the isotropic generalized diffusion tensor MRI (IGDTI) acquisition protocol to achieve an efficient orientationally averaged DW signal [3] with the following parameters: 16 linearly sampled $b$-values ranging from 2,540 to 14,700 s/mm$^2$ in 3 directions, 14 linearly sampled b-values ranging from 4,140 to 14,700 s/mm$^2$ in 4 directions, and 9 linearly sampled b-values ranging from 8,260 to 14,700 s/mm$^2$ in 6 directions, using the efficient gradient sampling schemes in Table 2 in [3]. This type of diffusion encoding increases the contrast given by local anisotropy and is not intended to measure the isotropic diffusion in the system. Additional diffusion parameters were gradient duration of $\delta = 4$ ms and diffusion time of $\Delta = 15$ ms.

The three 2D distributions of MD-$T_1$, MD-$T_2$, and $T_1$-$T_2$, were estimated, respectively, with the following data acquisition protocols (in conjunction with the *a priori* obtained 1D distributions as constraints): A 2D diffusion-$T_1$-weighted data set with 16 sampled combinations of inversion times and b-values within the aforementioned 1D acquisition range; a 2D D-$T_2$-weighted data set with 16 sampled combinations of echo times and b-values within the aforementioned 1D acquisition range; and a 2D $T_1$-$T_2$-weighted data set with 16 sampled combinations of inversion and echo times within the aforementioned 1D acquisition range.

The data were averaged 4 times to maintain high signal-to-noise ratio (SNR), which was always maintained above 100 (defined as the ratio between the average unattenuated signal intensity within a tissue region of interest, and the standard deviation of the signal intensity within the background). The sample temperature was set at 16.8°C.

### Immunohistochemistry

Primary antibodies used were antiglial fibrillary acidic protein (GFAP, mouse antihuman monoclonal antibody GA5 with bond heat-induced epitope retrieval, epitope retrieval time 10minutes, PA0026; Leica Biosystems, Wetzlar, Germany), amyloid precursor protein (APP, mouse antihuman monoclonal antibody clone 22c11, dilution 1:10, epitope retrieval time 10 minutes, MAB348; EMD Millipore, Burlington, MA), and abnormally phosphorylated tau (AT8; mouse anti-human monoclonal antibody, dilution 1:2000, Thermo Scientific, MN1020, HIER 1:10 for 10 min).

### References

[1] Benjamini, D. & Basser, P. Use of marginal distributions constrained optimization (madco) for accelerated 2d mri relaxometry and diffusometry. *Journal of Magnetic Resonance* **271**, 40–45 (2016). https://doi.org/10.1016/j.jmr.2016.08.004 .

[2] Benjamini, D. *et al.* Diffuse axonal injury has a characteristic multidimensional mri signature in the human brain. *Brain* **144**, 800–816 (2021). https://doi.org/10.1093/brain/awaa447 .

[3] Avram, A. V., Sarlls, J. E., Hutchinson, E. & Basser, P. J. Efficient experimental designs for isotropic generalized diffusion tensor MRI (IGDTI). *Magnetic Resonance in Medicine* **79** (1), 180–194 (2018) .
